# Genetic architecture, spatial heterogeneity, and the coevolutionary arms race between newts and snakes

**DOI:** 10.1101/2023.12.07.570693

**Authors:** Victoria Caudill, Peter L. Ralph

## Abstract

Coevolution between two species can lead to exaggerated phenotypes that vary in a correlated manner across space. However, the conditions under which we expect such spatially varying coevolutionary patterns in polygenic traits are not well-understood. We investigate the coevolutionary dynamics between two species undergoing reciprocal adaptation across space and time, using simulations inspired by the *Taricha* newt – *Thamnophis* garter snake system. One striking observation from this system is that newts in some areas carry much more tetrodotoxin than in other areas, and garter snakes that live near more toxic newts tend to be more resistant to this toxin, a correlation seen across several broad geographic areas. Furthermore, snakes seem to be “winning” the coevolutionary arms race, i.e., having a high level of resistance compared to local newt toxicity, despite substantial variation in both toxicity and resistance across the range. We explore how possible genetic architectures of the toxin and resistance traits would affect the coevolutionary dynamics by manipulating both mutation rate and effect size of mutations across many simulations. We find that coevolutionary dynamics alone were not sufficient in our simulations to produce the striking mosaic of levels of toxicity and resistance observed in nature, but simulations with ecological heterogeneity (in trait costliness or interaction rate) did produce such patterns. We also find that in simulations, newts tend to “win” across most combinations of genetic architectures, although the species with higher mutational genetic variance tends to have an advantage.

## 1 Introduction

Coevolution is the reciprocal adaptation of heritable traits between interacting species (Janzen 1980). These dynamic interactions between species shape patterns of adaptation, genetic diversity, and ecological dynamics. Coevolution shapes our world through mutualistic partnerships (such as pollinators and plants) (Janzen 1966), symbiotic dependencies (Thrall et al. 2007) and arms races (e.g., between host and pathogen or predator and prey) (Daugherty and Malik 2012). These multifarious interactions not only contribute to biodiversity, but also define species relationships, and accentuate the development of exaggerated traits. A better understanding of coevolution will help us to better craft conservation strategies, efficiently manage pest populations, and understand the dynamics that underpin stable ecosystems.

A coevolutionary interaction between species in which the individuals of one species benefit from the interaction and the individuals of the other is being harmed or killed and where phenotypes are continuously escalating is called an antagonistic coevolutionary arms race. For instance, a prey might evolve a defense mechanism and in response, its predator might respond by evolving a counter-mechanism, resulting in an ongoing cycle of adaptations and counter-adaptations (e.g., between herbivore resistance and plant defenses, Ehrlich and Raven 1964). Arms races between species have been observed in many species pairs such as *Taricha* newts and *Thamnophis* garter snakes (Brodie 2003), bacteria and phage (Bohannan and Lenski 2000), flax and flax rust (Dodds et al. 2006), and parsnip and parsnip webworm (Berenbaum, Zangerl, and Nitao 1986).

Antagonistic coevolution can be complex, especially when multiple interactions occur across space and time (Forde, Thompson, and Bohannan 2004). Current theory suggests that spatial structure facilitates coevolution by constraining phenotypes in local populations that differ across larger geographical areas (Gibert et al. 2013). Other theory has described how the spatial scales of patterns in host-parasite coevolution are determined by spatial movement and the nature of the coevolutionary interaction (Week and Bradburd 2023). An ambitious attempt to describe the spatial dynamics of coevolution, called the geographic mosaic theory, postulates that spatial structure and environmental heterogeneity creates coevolutionary “hot” and “cold” spots in the arms race, leading to a mosaic of selection pressures and a hypothesized dynamic process known as “trait remixing” (Thompson 2005). Recent studies have tried to infer the contribution of coevolution to speciation and diversification (Hembry, Yoder, and Goodman 2014), but there remains uncertainty about the degree to which coevolution is a primary mechanism for diversification of traits (Eaton 2008; Butler et al. 2009; Thompson 2009; Hembry, Yoder, and Goodman 2014; Parchman et al. 2016).

The well-studied predator-prey *Taricha* newt/*Thamnophis* garter snake system is relatively well-understood and features an intriguing spatial mosaic of trait variation over a wide geographical range (Brodie, Feldman, et al. 2005; Hanifin, Brodie, and Brodie 2008; Tseng 2011; Hague, Stokes, et al. 2020a; Reimche et al. 2020). On the Pacific Northwest coast of North America, various species of rough-skinned newt in the genus *Taricha* often contain tetrodotoxin that poisons predators. One of their predators, garter snakes in the genus *Thamnophis*, has developed resistance to this toxin. Levels of toxicity and resistance are highly correlated across the region: areas where newt toxicity is high, snake resistance is usually also high; and in areas where newt toxicity is low, snake resistance is usually also low. Furthermore, the snakes appear to be “winning” the coevolutionary arms race: snakes in each area seem to be able to eat the local newts with relatively little ill effect, no matter their toxicity (Hanifin, Brodie, and Brodie 2008; Feldman et al. 2010).

These two striking observations – that the coevolutionary outcome varies strongly across geog-raphy, and that the snakes nonetheless seem to be ahead of the newts – provide additional clues about the underlying biological basis of the interaction. Despite substantial work, the underlying cause of the observed geographic mosaic of hotspots and coldspots is still unknown. Such mosaics of coevolving traits are generally thought to be the result of heterogeneous ecological factors (e.g., resource availability and/or differences in community composition), nonadaptive forces (e.g., local genetic drift and population structure), or both (Brodie, Ridenhour, and Brodie 2002; Hague, Stokes, et al. 2020a; Thompson 2005). It would be of substantial interest to know the balance of these forces in this particular case. To explain the observation that snakes tend to be “winning”, Feldman et al. (2010) suggested that the availability of large-effect resistance alleles allowed the snakes a “potential ‘escape’ from the arms race”. This leaves us with two questions: *Is* heterogeneity in ecological factors required to explain the strikingly correlated maps presented in Hanifin, Brodie, and Brodie (2008) and Hague, Stokes, et al. (2020b), or can nonadaptive forces lead to a mosaic? And, *does* a less-polygenic architecture provide an advantage in this antagonistic coevolutionary arms race?

To answer these questions, as well as to better understand the dynamic interaction between spatial structure, genetic architecture, and coevolution, we conducted a simulation study, exploring a range of situations plausible for the *Taricha* newt/*Thamnophis* garter snake system. In addition to answering these specific questions, it is intriguing to consider other possible evolutionary outcomes, and what conditions made this outcome possible. For instance, it is easy to imagine alternate worlds in which snakes cannot eat newts (and survive), or in which snakes only eat newts in locations where newts are less toxic. The many determinants of the coevolutionary outcome include the strengths of various aspects of selection, and the genetic opportunity for adaptation. The genetic basis of these traits influences how the traits vary within populations and how they respond to selective pressures and environmental factors (Hoeksema and Forde 2008; Feldman et al. 2010), which can lead to different evolutionary outcomes. In snakes, there are three known gene mutations that lead to high levels of tetrodotoxin resistance (Feldman et al. 2012; McGlothlin et al. 2014). Each of these mutations alters ion channel functioning, and so decreases the ability of a snake carrying the mutation to crawl. The frequencies of each of these mutations vary over geographical space, and do not strongly correlate with local levels of newt toxicity (Geffeney et al. 2005). On the other hand, the genetic basis of newt toxicity is still unknown. A newt’s level of toxicity might be inducible and might be a result of environmental or bacterial factors. It is still unknown if it is heritable (Bucciarelli, Shaffer, et al. 2017; Bucciarelli, Alsalek, et al. 2022a; Vaelli et al. 2020). However, Bucciarelli, Shaffer, et al. (2017) showed that newts retain a base level of toxicity even when kept in lab conditions, and that young newts who are more toxic take longer to develop.

To better understand the process of adaptive evolution across geographical space, this paper asks three main questions: (1) How does the genetic architecture of the traits in newts and snake affect how they coevolve? (2) Under what situations do we get spatial patterns of correlated traits in newts and snakes as we see in the real world? (3) How fast does coevolution increase resistance and toxicity in these organisms with different combinations of genetic architectures? In particular, we compare different levels of genetic variance and polygenicity using individual-based simulations of continuous geographic space. The results complement field observations by describing situations that are consistent with empirical observations, and exploring other possible outcomes.

## 2 Methods

To explore these questions, we ran spatial individual-based, forward in time simulations with SLiM (version 3, Haller and Messer 2019). The simulation had two species that we call: “newts” and “snakes”, each with a quantitative trait for toxicity and resistance, respectively. The simulated range was a large rectangular map that spans 35 units in width and 140 units in length (long rectangular shape similar to the west coast, larger simulations hampered the speed of the simulation). The demographic details of this model were motivated by our current understanding of the coevolutionary interactions of the rough-skinned newt (*Taricha granulosa*) and the garter snake (*Thamnophis sirtalis*). These simulations are a simplistic representation of the system’s ecological and demographic complexities, but aim to explore important aspects of the possible interaction dynamics.

### The demographic model

Each simulated individual newt or snake was hermaphroditic and had a genome of 10^8^ base pairs, on which phenotype-affecting mutations occurred at a rate *µ* base pair per generation. (Although real newts and snakes of these species are not hermaphroditic, this should not significantly affect the dynamics). Each new mutation had an “effect size” drawn from a Normal distribution with a mean of zero and a standard deviation of *σ* (the values of *µ* and *σ* differed between simulations runs, as described below). Demographic parameters were chosen that both newts and snakes live for about 4 generations, on average. Newts and snakes had phenotypes we call “toxicity” and “resistance”, respectively, that are determined by genotypes in a way specified below. Each species also had fixed values for recombination rate (10*^—^*^8^ crossovers per bp per generation) and for parameters controlling offspring dispersal and mate selection. The simulations used SLiM’s “non-Wright-Fisher” population model (Haller and Messer 2019) with overlapping generations and fluctuating population sizes, with population dynamics described below. (We used SLiM version 3.7.1 with different “populations” for each species; since then in version 4.0 direct support for multiple species was added (Haller and Messer 2023).)

Every generation, each newt finds a nearby newt with whom to mate and produce offspring. A newt at spatial location *x* with *k* neighbors at locations *x*_1_, . . ., *x_k_* would choose neighbor *i*as a mate with probability

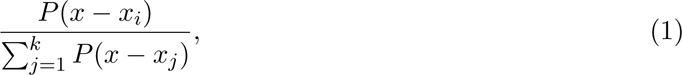

where *P* () is the density of a 2-D Normal distribution with standard deviation of 1 unit. The number of offspring they produce is Poisson distributed with a mean of 1/4, and each offspring thus produced would disperse to a random location whose displacement relative to the parent was drawn from a Normal distribution with a mean of zero and a standard deviation of 1 unit. Local newt population density was computed by smoothing using a Normal kernel: local density at location *x* is

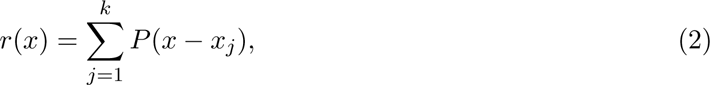

where *x*_1_*,. .., x_j_* are the locations of nearby newts (up to a maximum distance of 3 units).

Higher phenotypes were more costly than lower phenotypes. The probability that a newt at location *x* survives to the next time step if they have phenotype *z* is

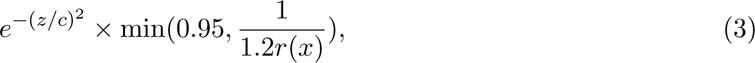

where *c* is a parameter controlling the costliness of the phenotype (so that fitness decreases as the phenotype increases). The probability of survival decreases with density, due to competition between individuals of the same species. Consequently, this leads to higher mortality in areas of higher densities. The parameters are chosen so that the rough equilibrium density is one individual per unit area. Unless otherwise stated, the value of *c* was set to 100.

Snake reproduction, offspring dispersal, and survival used the same dynamics as the newts. Furthermore, at each time step, each nearby newt-snake pair could “encounter” each other. To do this, we iterated over all snakes in random order; choosing for each snake a set of nearby newts and resolving these encounters before moving to the next snake. The probability that a snake “encounters” a given newt at distance *D* is

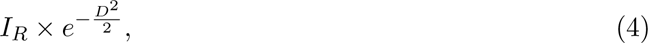

where *I_R_* is the baseline interaction rate for nearby individuals (set to 0.05 unless otherwise noted), and that individuals that were closer were more likely to interact than individuals further apart. When a snake and a newt encounter each other, the outcome of the interaction depends on the difference between their phenotypes: if *L* is the newts toxicity minus the snake’s resistance, then the probability that the snake survives and the newt dies is

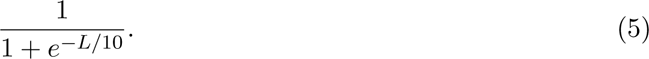

The form is chosen so that phenotype differences must change by around 10 units to substantially affect the interaction. If the snake does not eat the newt, the snake dies and the newt survives. For a newt to have a good chance of surviving an interaction with a snake, their level of toxicity needs to be greater than the snake’s level of resistance. For each newt a snake consumes, the snake receives a “fitness bonus” of 0.1, which is added to the probability of surviving to the next time step (however, increases past 1.0 have no effect).

We used coalescent simulations produced by msprime (Kelleher, Etheridge, and McVean 2016) to generate initial genetic variation. These simulations had an effective population size of 10,000 and a recombination rate of 10*^—^*^8^. Mutations’ effect sizes were drawn from the same distribution as within SLiM, and added to the resulting tree sequence using pyslim (Rodrigues et al. 2023) (these had no effect on the coalescent simulation). To initialize each SLiM simulation, 300 individuals for each species were uniformly distributed across space.

### Genetic Architecture

A major goal of our study is to describe how genetic architecture affects this coevolving system. By “genetic architecture” we mean the combination of mutation rate (*µ*) and the standard deviation of mutation effect size (*σ*, which we refer to as “effect size”). We explored a range of values for mutation rate and effect size to span from a highly polygenic to an oligogenic model (Table 1). Many evolutionary dynamics depend primarily on mutational variance, which is

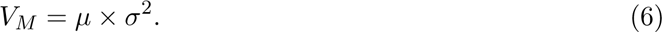

We model the phenotypes for each species as exponentials of additive genetic traits. Concretely, if the sum of the effects of all mutations carried by an individual newt is *G*, then that newt’s phenotype (i.e., toxicity) is *e^G/^*^10^. Snake resistance is determined in exactly the same way. The exponential transform is used here because toxicity and resistance are non-negative, and because then mutations have multiplicative effects (i.e., increase or decrease the phenotypes by percentages).

### Simulation Experiments

We ran four replicate simulations at each distinct set of parameter values, across all experiments. Table 1 shows the parameters used in three experiments that test how mutational variance impact the evolution of newt and snake phenotypes through different combinations of mutation rate and mutation effect size. Our first experiment set (Experiment 1) varies both mutation rates and mutation effect sizes. In the second experiment, mutation rate was for both species in each simulation, but allows the species to have different mutation effect sizes (and hence mutational variance). In the third experiment, mutational variance (*V_M_*) was the same for newts and snakes in each simulation, although polygenicity could be different (varying mutation rate and mutation effect size).

Experiment 1 ran simulations with all possible combinations of the four genetic architectures labeled “1a” to “4a” in Table 1. Since there are sixteen possible combinations (e.g., snakes had 1a and newts had 3a) and we ran four replicates per combination, there was a total of 64 simulations. Across these genetic architectures, mutation rate that ranged from 10*^—^*^8^ to 10*^—^*^11^, and the standard deviation of mutation effect sizes (*σ*) ranged from 0.005 to 0.05. The genetic architectures are arranged so that mutational variance increases with label (see rightmost column in Table 1.

In Experiment 2, we study the effects of mutational variance. To do this, we spanned the same range of parameters as in Experiment 1, in each simulation both newts and snakes have the same mutation rate (*µ*), but have (possibly) different mutation effect sizes (*σ*), and hence different mutational variances (*V_M_* = *µσ*^2^). Note that in Table 1, genetic architectures with the same mutation rate are grouped together in shaded groups of four rows, and that architectures in the same group have the same numbered portion of their label. So, in each simulation as a part of this experiment, newts and snakes were each assigned a genetic architecture from the same shaded group: this led to sixteen combinations within each of the four mutation rate groups, and hence 256 simulations across 64 distinct combinations of genetic architectures.

In Experiment 3 we matched mutational variance between the species, and looked for the effects of polygenicity. This was structured similarly to Experiment 2: shaded groups in Table 1 now have the same *V_M_*, and we ran simulations with each of the sixty-four possible pairs of genetic architectures for which the two are drawn from the same group (with replicates, 256 total simulations). For instance, in one simulation, newts had genetic architecture with *µ* = 10*^—^*^8^ and *σ* = 0.0158 “1g” and snakes had genetic architecture with *µ* = 10*^—^*^11^ and *σ* = 0.5 “4g”; these two have the same mutational variance (*V_M_* = 2.5 *×* 10*^—^*^12^), but different polygenicity (newts have a high rate of small mutations; snakes have a low rate of large mutations). (Also note that the set of genetic architectures in Experiment 3 is the same as in Experiment 2; what differs between the experiments is which pairs are assigned to newts and snakes.)

### The contribution of coevolution to phenotype change

To assess the strength of coevolution on phenotype change within the simulations (as opposed to genetic drift), we ran additional simulations after modifying the original model. We separately made two important changes, making either (1) the trait (toxicity or resistance) not heritable, and (2) the outcome of the interaction random (instead of dependent on phenotype). For (1), every new individual had a phenotype not determined by their genetics but instead as *e^G/^*^10^ with *G* chosen independently from a Normal distribution with mean set to 3 and a standard deviation set at 2 (creating a phenotype near 1). For (2), we made the outcome of each snake-newt interaction depend on the result of a fair coin flip instead of the difference in phenotype: with 50% probability the snake eats a newt, otherwise the snake dies.

**Table 1:** A summary of all parameter sets used in simulations. Within each of the three experiments, newts and snakes were assigned all combinations of genetic architectures (i.e., rows of this table) for which both rows are in the same group (colored blocks). Parameter sets are labeled (second column) by an identifier so that two parameter sets are in the same group if they have the same letter in their label.

| Experiment | Genetic Arch.<br>(Newt/Snake) | Mutation<br>Rate ($\mu$) | Mutation Effect<br>Size ($\sigma$) | Mutational<br>Variance |
| --- | --- | --- | --- | --- |
| 1: Low to High<br>Mutational Variance<br>with changing<br>mutation rate | 1a | $10^{-8}$ | 0.005 | $2.5 \times 10^{-13}$ |
| | 2a | $10^{-9}$ | 0.05 | $2.5 \times 10^{-12}$ |
| | 3a | $10^{-10}$ | 0.5 | $2.5 \times 10^{-11}$ |
| | 4a | $10^{-11}$ | 5.0 | $2.5 \times 10^{-10}$ |
| 2: Low to High<br>Mutational Variance<br>(same mutation<br>rate) How does<br>increasing<br>mutational variance | 1b | $10^{-8}$ | 0.005 | $2.5 \times 10^{-13}$ |
| | 2b | $10^{-8}$ | 0.0158 | $2.5 \times 10^{-12}$ |
| | 3b | $10^{-8}$ | 0.05 | $2.5 \times 10^{-11}$ |
| | 4b | $10^{-8}$ | 0.158 | $2.5 \times 10^{-10}$ |
| | 1c | $10^{-9}$ | 0.0158 | $2.5 \times 10^{-13}$ |
| | 2c | $10^{-9}$ | 0.05 | $2.5 \times 10^{-12}$ |
| | 3c | $10^{-9}$ | 0.158 | $2.5 \times 10^{-11}$ |
| | 4c | $10^{-9}$ | 0.5 | $2.5 \times 10^{-10}$ |
| | 1d | $10^{-10}$ | 0.05 | $2.5 \times 10^{-13}$ |
| | 2d | $10^{-10}$ | 0.158 | $2.5 \times 10^{-12}$ |
| | 3d | $10^{-10}$ | 0.5 | $2.5 \times 10^{-11}$ |
| | 4d | $10^{-10}$ | 1.58 | $2.5 \times 10^{-10}$ |
| | 1e | $10^{-11}$ | 0.158 | $2.5 \times 10^{-13}$ |
| | 2e | $10^{-11}$ | 0.5 | $2.5 \times 10^{-12}$ |
| | 3e | $10^{-11}$ | 1.58 | $2.5 \times 10^{-11}$ |
| | 4e | $10^{-11}$ | 5.0 | $2.5 \times 10^{-10}$ |
| 3: Same Mutation<br>Variance How does<br>effects of<br>polygenicity effect<br>newt and snake<br>phenotype? | 1f | $10^{-8}$ | 0.005 | $2.5 \times 10^{-13}$ |
| | 2f | $10^{-9}$ | 0.0158 | $2.5 \times 10^{-13}$ |
| | 3f | $10^{-10}$ | 0.05 | $2.5 \times 10^{-13}$ |
| | 4f | $10^{-11}$ | 0.158 | $2.5 \times 10^{-13}$ |
| | 1g | $10^{-8}$ | 0.0158 | $2.5 \times 10^{-12}$ |
| | 2g | $10^{-9}$ | 0.05 | $2.5 \times 10^{-12}$ |
| | 3g | $10^{-10}$ | 0.158 | $2.5 \times 10^{-12}$ |
| | 4g | $10^{-11}$ | 0.5 | $2.5 \times 10^{-12}$ |
| | 1h | $10^{-8}$ | 0.05 | $2.5 \times 10^{-11}$ |
| | 2h | $10^{-9}$ | 0.158 | $2.5 \times 10^{-11}$ |
| | 3h | $10^{-10}$ | 0.5 | $2.5 \times 10^{-11}$ |
| | 4h | $10^{-11}$ | 1.58 | $2.5 \times 10^{-11}$ |
| | 1i | $10^{-8}$ | 0.158 | $2.5 \times 10^{-10}$ |
| | 2i | $10^{-9}$ | 0.5 | $2.5 \times 10^{-10}$ |
| | 3i | $10^{-10}$ | 1.58 | $2.5 \times 10^{-10}$ |
| | 4i | $10^{-11}$ | 5.0 | $2.5 \times 10^{-10}$ |

To measure the speed of coevolution, we identified the “final mean phenotype” as the average mean phenotype over the last 1000 time steps, and the “equilibrium time” as the first time mean phenotype reached 98% of the final mean phenotype. We then reported the speed as the difference in mean phenotype between the equilibrium time and time step 100, divided by the number of elapsed time steps.

### Heterogeneous landscapes

We also conducted simulations where some parameters changed across geographical space. In these simulations, we used the genetic architectures from Experiment 1 (see Table 1) to see how newt and snake phenotypes changed when the costliness of the phenotypes or the interaction rate varied across space. In each, the relevant parameter varied across the map in a linear gradient. Each individual’s location was then used to determine the correct value for that individual.

To simulate geographical variation in phenotype cost, the parameter *c* of equation (3) decreased linearly across the map, from *c* = 50 in the south to *c* = 250 in the north. So, individuals that lived in the top portion of the map had a larger fitness penalty for having a high phenotype than did individuals living in the bottom portion of the map. We ran simulations in which there was geographical variation in cost for both newts and snakes, as well as simulations in which cost varied for only one species. The fitness cost ranged from 50 to 250 and impacted the probability of snake and newt survival (equation 3).

To simulate geographical variation in interaction rate, each snake’s location was used to determine the interaction rate, i.e., the parameter *I_R_* from equation (4), which is the base probability with which a snake would encounter a nearby newt. This map had newts and snakes interacting more at the bottom of the map than at the top of the map: *I_R_* ranged from 0.01 to 0.1 linearly with north-south position.

### Data collection

We used code in SLiM to collect data on newt and snake phenotypes across the entire geographical area. In particular, we collected local newt and snake mean phenotypes by using SLiM’s summarizeIndividuals() function to divide the map up into a 5 *×* 20 grid and calculating the local mean phenotypes within each grid cell. The local mean phenotypes were then used to calculate spatial correlations between newt and snake phenotypes.

### Data availability

The simulation and analysis code is accessible through github (github link to be added).

## 3 Results

### Newt and snake Evolution

We first evaluated under which situations where phenotypes were actually coevolving. If coevolution was occurring, we anticipated an increase in newt and snake average phenotypes over time. Figure 1 shows three common outcomes of the simulation, with mean phenotypes for newts (red) and snakes (blue): fast evolution, slow evolution, and no evolution. Simulation with relatively fast evolution had the average newt and snake phenotype rising rapidly and reaching a steady-state point before 10,000 generations, while relatively slow simulations took longer than 20,000 generations to equilibrate as “slow”. In simulations with no evolution, the average phenotype of one or both species’ changed very little throughout the simulation.

**Figure 1:**
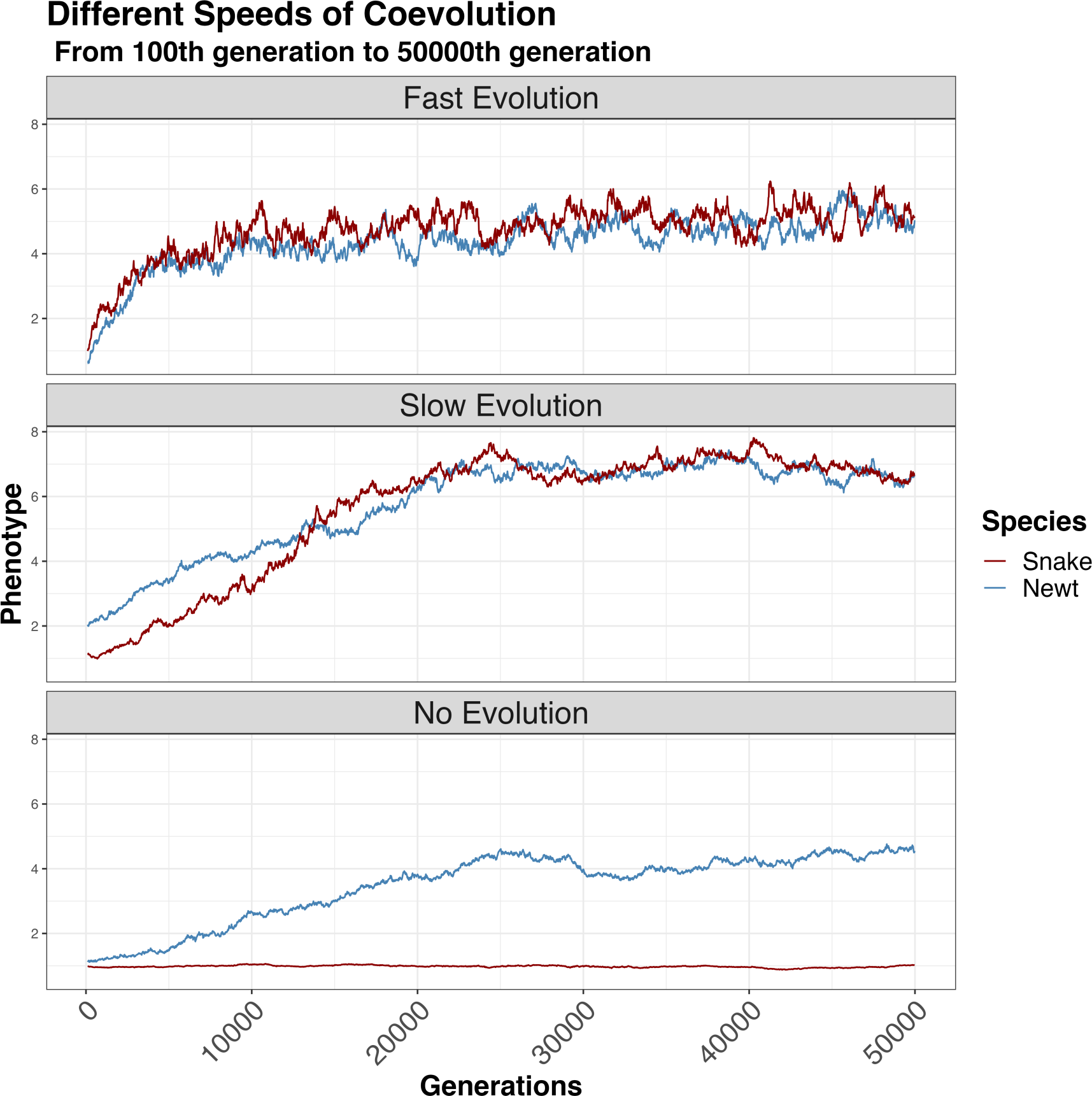
Mean phenotype dynamics over time in simulations showing three speeds of evolution; fast, slow, and no. In the top panel (fast evolution), newt (red) and snake (blue) average phenotypes go up quickly, reaching an equilibrium at around 10,000 generations. In the middle panel (slow evolution), newt and snake phenotypes rise more slowly, reaching equilibrium at around 20,000 generations. In cases of “no evolution”, at least one species’ mean phenotype remains flat, in this example (bottom panel), the mean newt phenotype remains flat.

Figure 2 shows the average change phenotypes (averaged across both species) over the course of the simulation for all genetic architecture combinations from Experiment 1. When the genetic architecture of either species had low mutational variance (this was genetic architecture 1a, with lowest *V_M_*), there was no coevolution: it’s mean phenotype did not increase (or decrease), regardless of the other species’ genetic architecture. However, even when one species could not evolve, the other species’ phenotype still might, due to genetic drift (e.g., snake phenotype increased when newt phenotype could not increase in Figure 1C). The strongest effect of coevolution – i.e., the largest change in both species’ phenotypes occurred when both had intermediate mutational variance and polygenicity (genetic architectures 2a and 3a). A species with the final genetic architecture (4a), which had the largest amount of mutational variance (primarily from large-effect mutations) still was able to coevolve, but showed a lower mean phenotype change.

**Figure 2:**
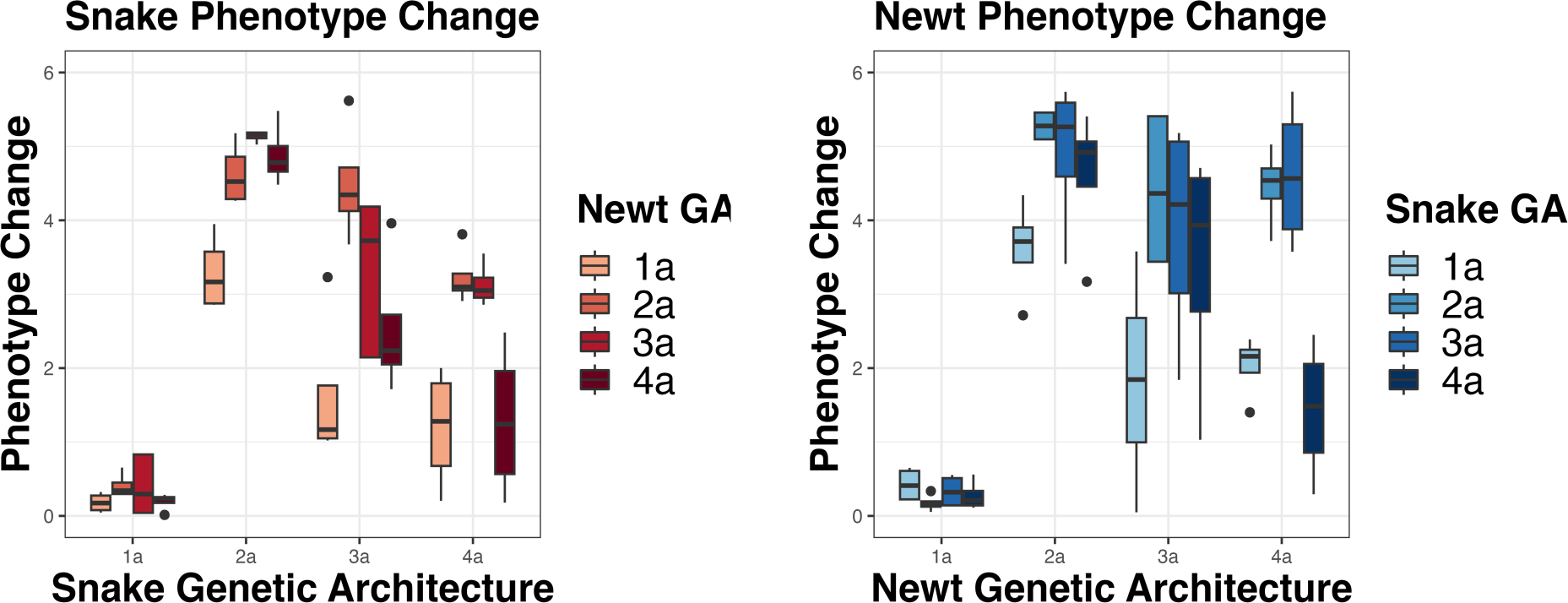
Mean phenotypes for each combination of genetic architectures in Experiment 1. Each box plot shows the range of mean phenotypes observed across time (i.e., the entire simulation) and simulation replicate for one combination of genetic architectures. The *x*-axis represented the genetic architecture of snake (left) or newt (right), and color represents the genetic architecture of the opposite species. Note that when either species has genetic architecture 1a (with lowest genetic variance), that species has consistently low phenotype; otherwise, their mean phenotype depends on the genetic architecture of the other species.

It is interesting to note that the change in newt and snake mean phenotype is not symmetrical with reciprocal genetic architecture combinations. For example, when both newts and snakes had genetic architecture 2a the change in newt phenotype was larger than the change in snake phenotype. There was also a relationship between phenotype and population size (see Supplementary Figures S1 and S2): species with higher average phenotype tended to have larger population sizes, as one might expect if that species were “winning” the coevolutionary arms race. (Even though we saw newt and snake population sizes fluctuate we did not observe extinction.) In simulations without the lowest mutation variance (genetic architecture 1a), newt and snake mean phenotypes seemed to be coevolving. However, this pattern could in principle be caused by drift instead of coevolution. To exclude this possibility, we next modified key components of the simulation to examine what was driving the change in phenotype.

### How much is coevolution driving phenotype change?

To verify that phenotypic changes were the result of coevolution, we altered both heritability and (separately) the snake-newt interaction while keeping everything else as similar as possible, so we could see what phenotypic changes were expected in the absence of coevolutionary forces (see Methods for details). If genetic drift were responsible for the observed increase in phenotype, we would see similar increases in phenotype in these simulations in which coevolution was not possible.

Figure 3 shows the average change in phenotype in these simulations, averaged across both species, for different genetic architecture combinations across generations 100 to 50,000, compared to the previously described coevolutionary simulations shown in Figure 1. For clarity, we do not show simulations in which either species had low mutational variance (i.e., genetic architecture 1a). As expected, we saw an increase in newt and snake phenotype only in the standard simulations (orange boxplots), and not when either heritability or dependence of the interaction on phenotype was removed (other boxes). In simulations where the interaction does not depend on phenotype (green boxplots), phenotypes decreased. In these, the phenotype provided no benefit, so the mean phenotype dropped to a level determined by mutation-selection balance (where “selection” is due to the costliness of the phenotype). Simulations in which the trait was not heritable (barely visible, grey boxplots) had no change in newt and snake phenotype for all genetic architectures, as expected. (However, these simulations still provide a meaningful control from levels of spatial correlations expected in the absence of coevolution.)

**Figure 3:**
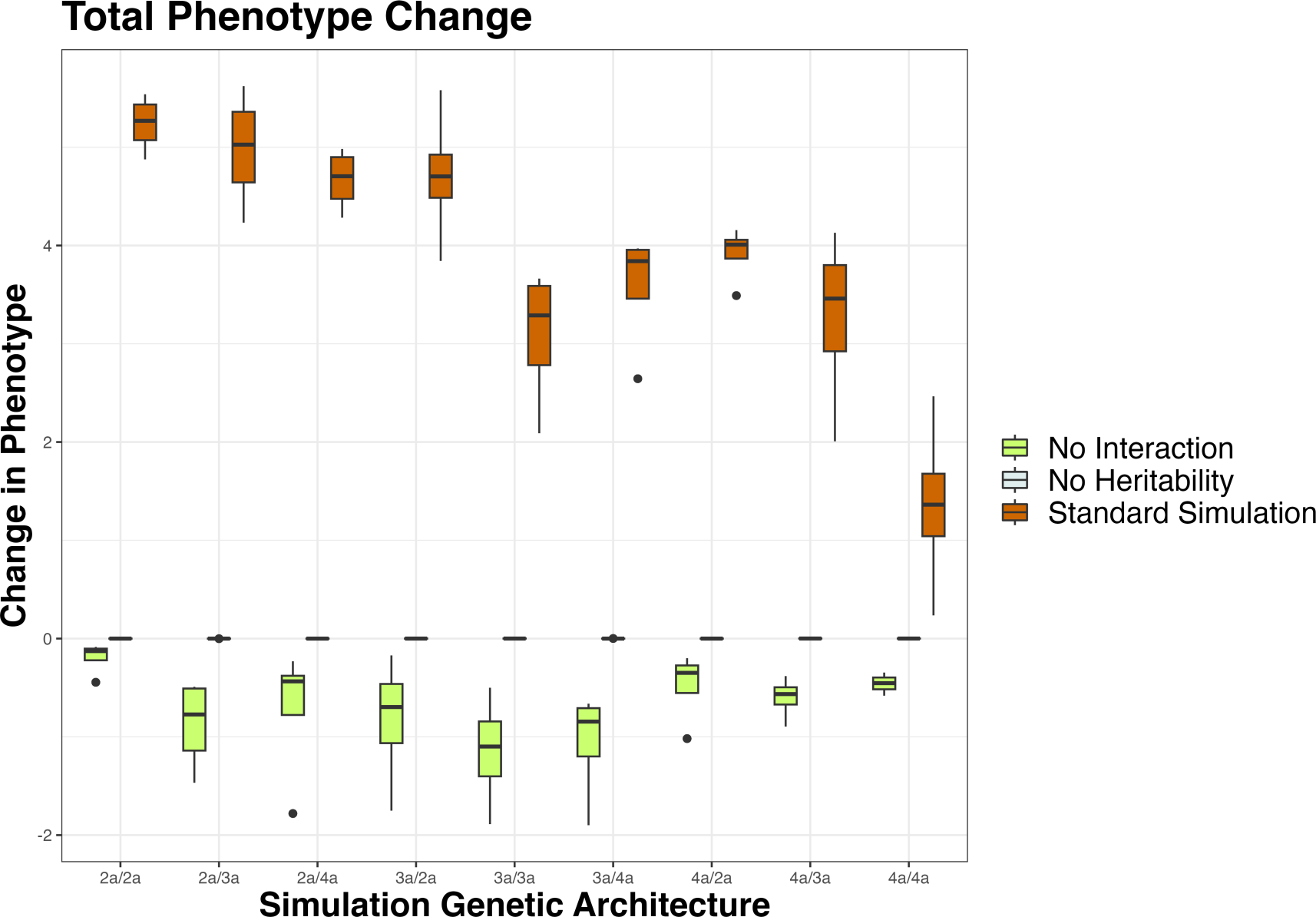
Newt and snake mean phenotypes (averaged together), comparing standard simulations to simulations without heritability or random interaction outcomes (see Methods). In the standard simulations (orange) the mean phenotype for newts and snakes increases. When there is no heritability (grey), phenotypes remain close to zero. When the newt-snake interaction outcome is random (not based on phenotype) the average newt and snake phenotypes decreased (green).

### Spatially Heterogeneous Landscapes

In the real world, newt and snake resistance are correlated across a broad geographical region (Hanifin, Brodie, and Brodie 2008; Reimche et al. 2020) in locations where newts are very toxic, snakes tend to be very resistant to their toxin, and vice-versa. The empirical (product-moment) correlation between the toxicity and resistance values reported in Hanifin, Brodie, and Brodie (2008) is *r* = 0.77. Do our simulations recapitulate this spatial correlation?

The answer, so far, is “no”: Figure 5 shows that no genetic architecture combination had a comparable degree of spatial correlation. (To obtain a roughly equivalent measure from our simulations, we split the entire area into 100 smaller squares (in a 5*×*20 grid), calculated local mean phenotypes of each species in the squares (Fig. 4), and computed the correlation between these two vectors of local mean phenotypes; see Figure 4 for an example.) Although local fluctuations driven by coevolution could in principle have created spatial correlations (Thompson 2005), it appears that such fluctuations do not appear strongly, at least across this range of parameter values in a homogeneous landscape.

**Figure 4:**
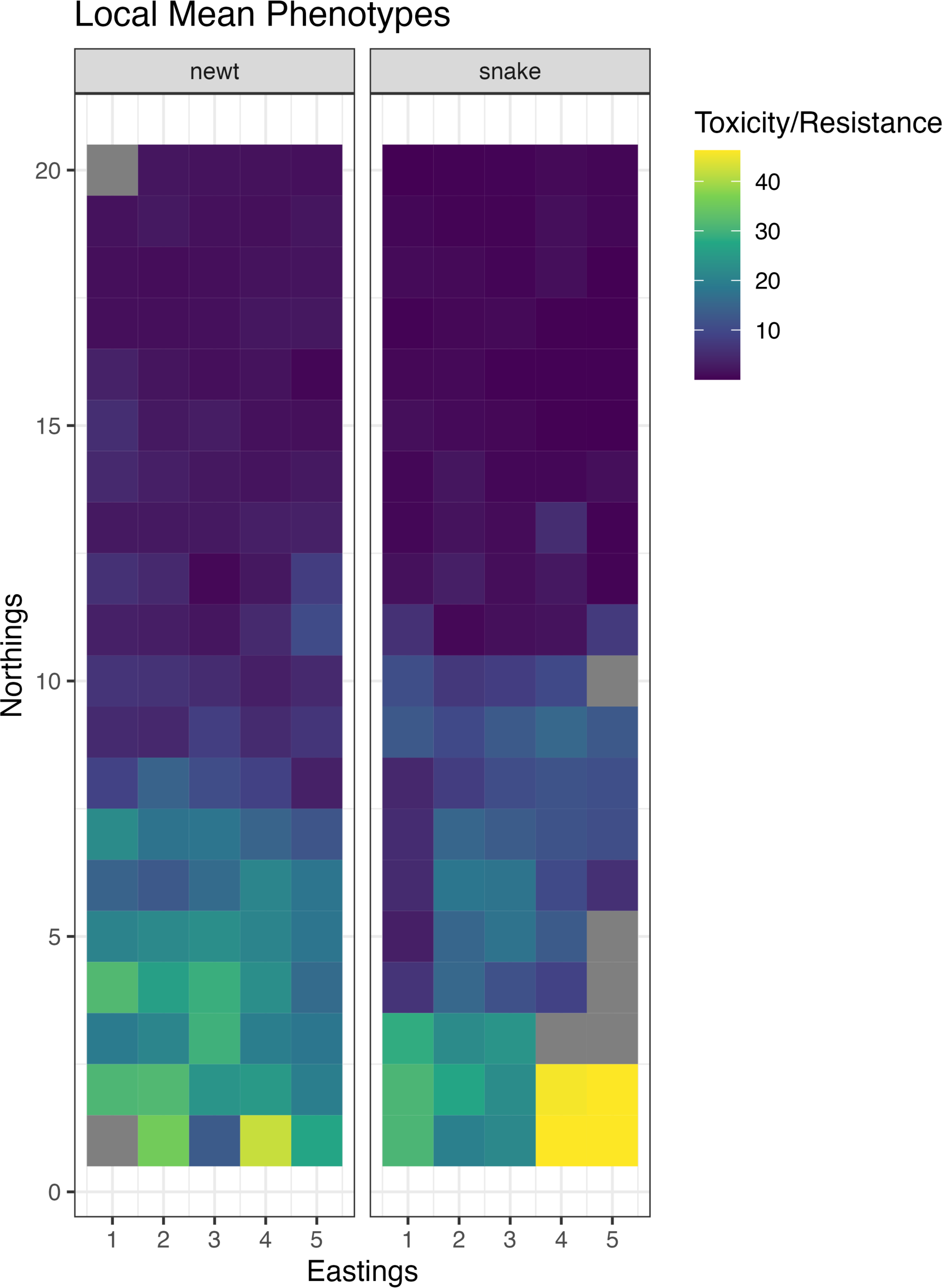
Local newt and snake mean phenotypes across the geographical area at generation 44501in a simulation in which costliness of the phenotype was a spatial gradient (higher costliness at the top of the map). The genetic architectures used were snake: 4a and newt: 3a; other combinations are roughly similar. Each box shows the mean ph^3^e^3^notype of individuals living within that grid cell. Brighter colors represent higher levels of toxicity or resistance.

**Figure 5:**
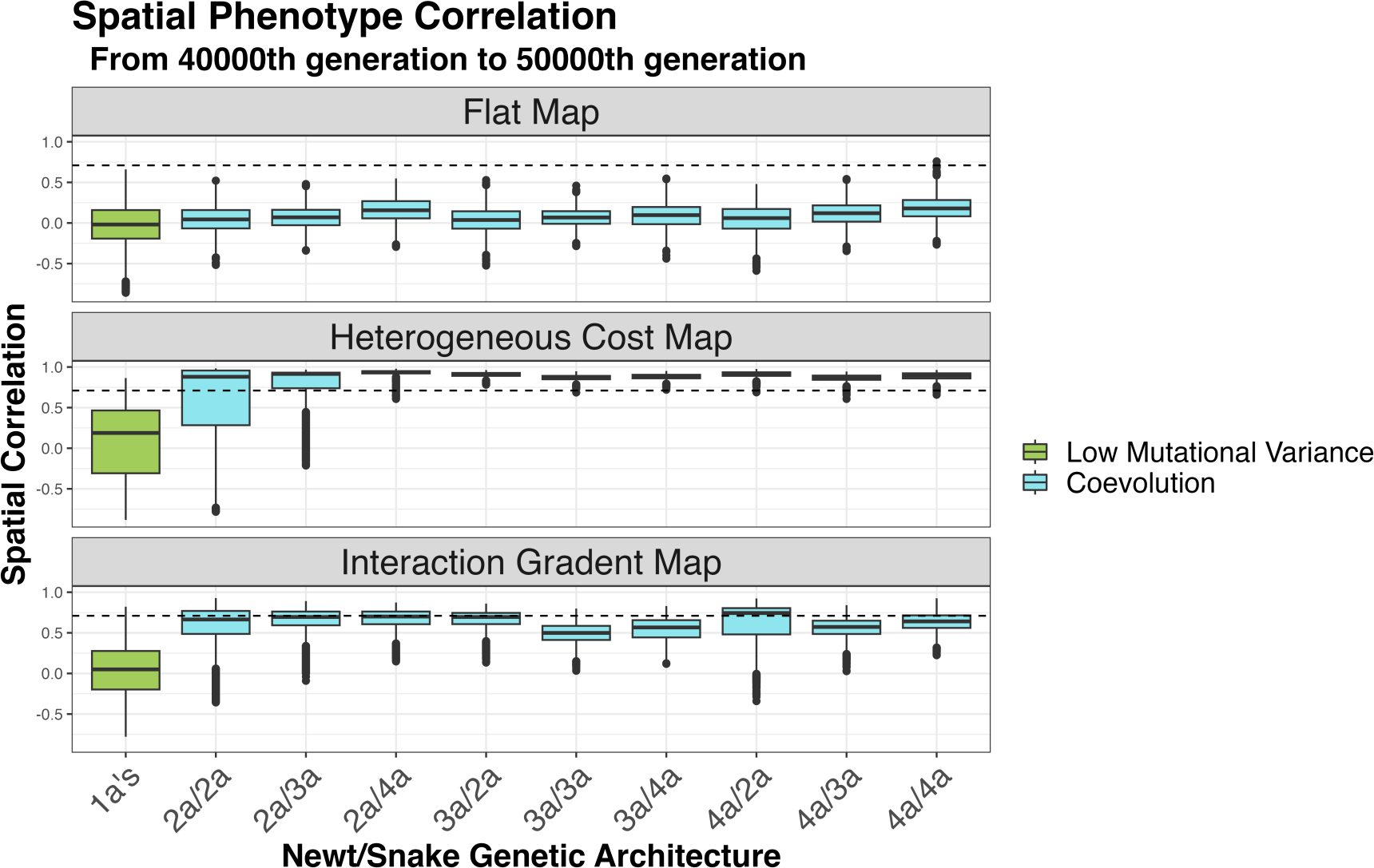
Spatial newt and snake phenotype correlations for all genetic architecture combinations, for three types of map. In each plot, the leftmost (green) boxplot displays correlations across all combinations of genetic architectures containing the lowest genetic variance (1a); other boxes (blue) show the range of spatial correlations across replicates and time steps. The dashed line shows the empirical newt and snake spatial correlation (reported by Hanifin, Brodie, and Brodie 2008). When there is no spatial heterogeneity in the simulation (top), there is little spatial correlation. Higher spatial correlations occur when newts and snake coevolve on a map with heterogeneity in phenotype costliness (middle) or interaction rate (bottom). Correlations are similar across all genetic architecture combinations with high enough mutational variance.

However, the landscape inhabited by the real newts and snakes is not constant. It seems very plausible that the real environment is heterogeneous – for instance, (Reimche et al. 2020) found that an elevation gradient in the Sierra Nevadas was correlated to levels of toxicity and resistance in a sister species of newts and snakes. Furthermore, variation in population density, predator pressure, and/or habitat could easily lead to varying rates of interaction between newts and snakes across the landscape. So, we also ran simulations in heterogeneous landscapes. We changed, in separate simulations, the costliness of the phenotype and the rate of interaction between newts and snakes across the map. When the costliness of the phenotype varied across space, phenotypes were even more strongly correlated than in the empirical data (mean *r* = 0.84, median *r* = 0.90, empirical *r* = 0.77). To make sure that this correlation caused by coevolution and not changing cost alone, we re-ran this simulation without a phenotype-based interaction (as described above) and saw no correlations (see Supplementary Figure S3). Simulations in which only one species’ costliness varied across the map had lower but still strong correlations (snakes vary: mean *r* = 0.47; median r = 0.49; newts vary: mean *r* = 0.57, median r = 0.70; see Supplementary Figure S3). In addition to having high correlations between phenotypes, all simulations with varying parameters across the map had a wide range of phenotypic values, as seen in the empirical data (as in Figure 4).

We also saw high correlations in simulations where the interaction rate varied across the map (mean *r* = 0.60). Varying interaction rate led to instances where local extinction and subsequent recolonizations occurred in small sections of the map. Adjusting the range of interaction rates or perhaps other parameters would likely increase correlations, but we did not explore parameter space further.

### Effects of Genetic Architecture on Coevolution

In Figure 2, we saw that the coevolutionary equilibrium values of newt and snake mean phenotypes differed substantially depending on the genetic architecture (and, interestingly, in an asymmetric way). However, we saw in Figure 5 that genetic architecture did not affect the spatial correlation of newt and snake phenotype. How else does genetic architecture affect the coevolutionary dynamics? And, does it affect which species is “winning”? Our simulations in Experiment 1 had genetic architectures that simultaneously varied mutational variance and polygenicity, so to disentangle these two effects, we ran two additional sets of simulations in which we (Experiment 2) constrained newts and snakes to have the same mutation rate (*µ*) but differing effect size (*σ*) and hence differing mutational variance (*V_M_* from equation (6)); and (Experiment 3) constrained newts and snakes to have the same mutational variance but different combinations of mutation rate and effect size.

We saw that the speed of coevolution depends more on mutational variance than it does on polygenicity for both newts (red) and snakes (blue) (Figure 6). When mutational variance was high the speed of coevolution was fast, essentially taking less time for the simulation to reach a steady-state. When mutational variance was lower, it took longer for the simulation to reach a steady-state.

**Figure 6:**
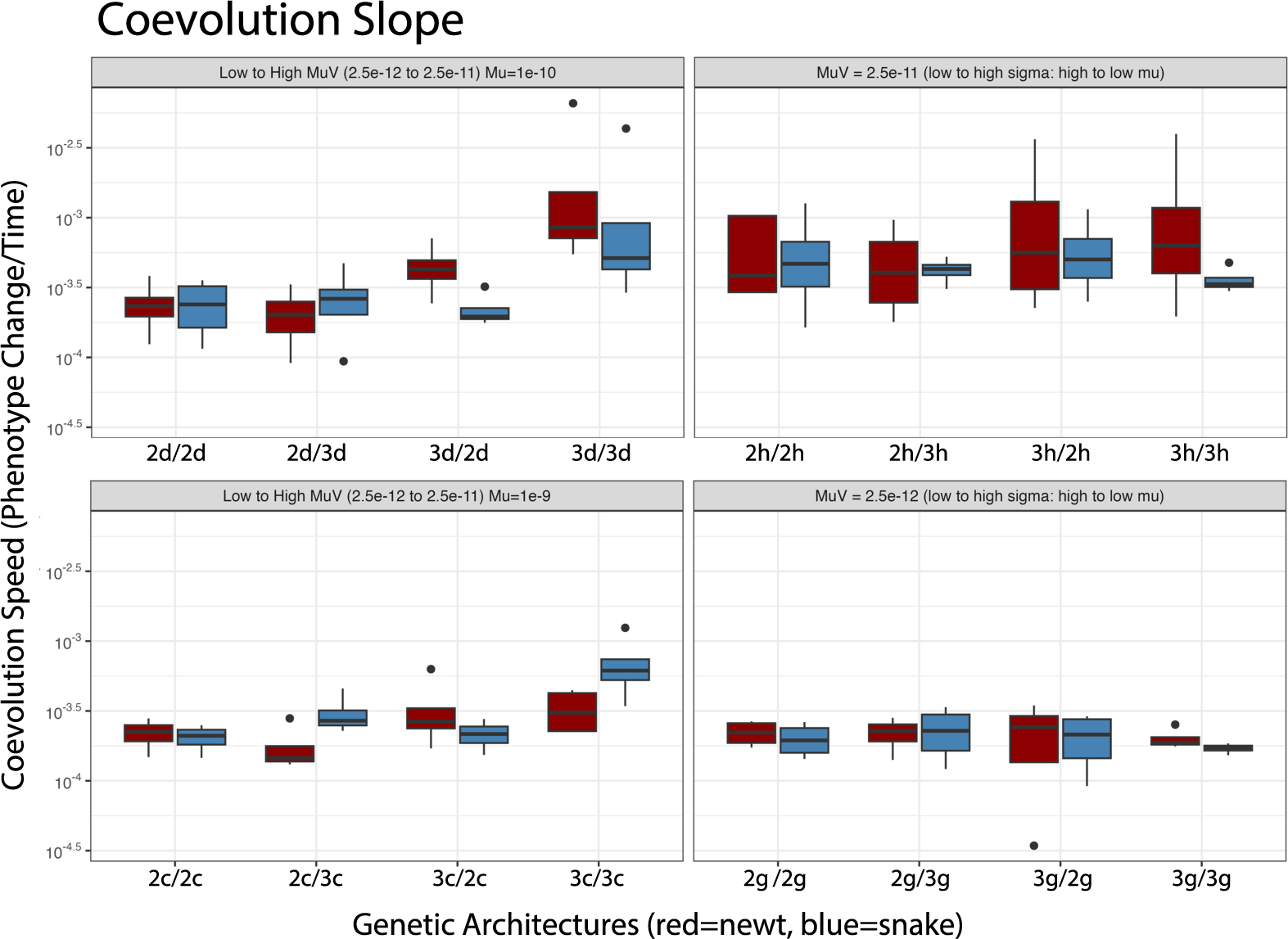
The speed of coevolution depends on mutational variance and not just the mutation rate or mutation effect size. Each boxplot shows the range of coevolution speed (see Methods) across simulation replicates at that combination of genetic architectures. **Left** plots show combinations in which mutation rate is matched between species, arranged so that mutational variance increases moving left to right. **Right** plots show combinations in which mutational variance is matched between species, arranged so that effect size (*σ*) increases and mutation rate (*µ*) decreases moving left to right. Note that the combination 3d/3d has the same amount of genetic variation as all combinations in the top-right panel (2h/2h, 2h/3h, 3h/2h, and 3h/3h) and that the combination 2c/2c has the same amount of genetic variation as those in the bottom-right panel.

Does it make sense to say that one species is “winning”? If one species has a higher mean phenotype than the other, then that species will more often “win” in encounters between the two species. Figure 7 shows mean phenotype differences between the species across the genetic architecture combinations of Experiment 1 (excluding those combinations with architecture 1a, as usual). Here, we see that population size differences and mean phenotype differences track each other (also see in Supplementary Figure S2). We also see that newts more often have the higher mean phenotype – apparently, the asymmetry in their ecological interaction gives newts the coevolutionary edge. However, in at least one situation of Figure 7, the snakes (on average) had a larger mean phenotype and population size.

**Figure 7:**
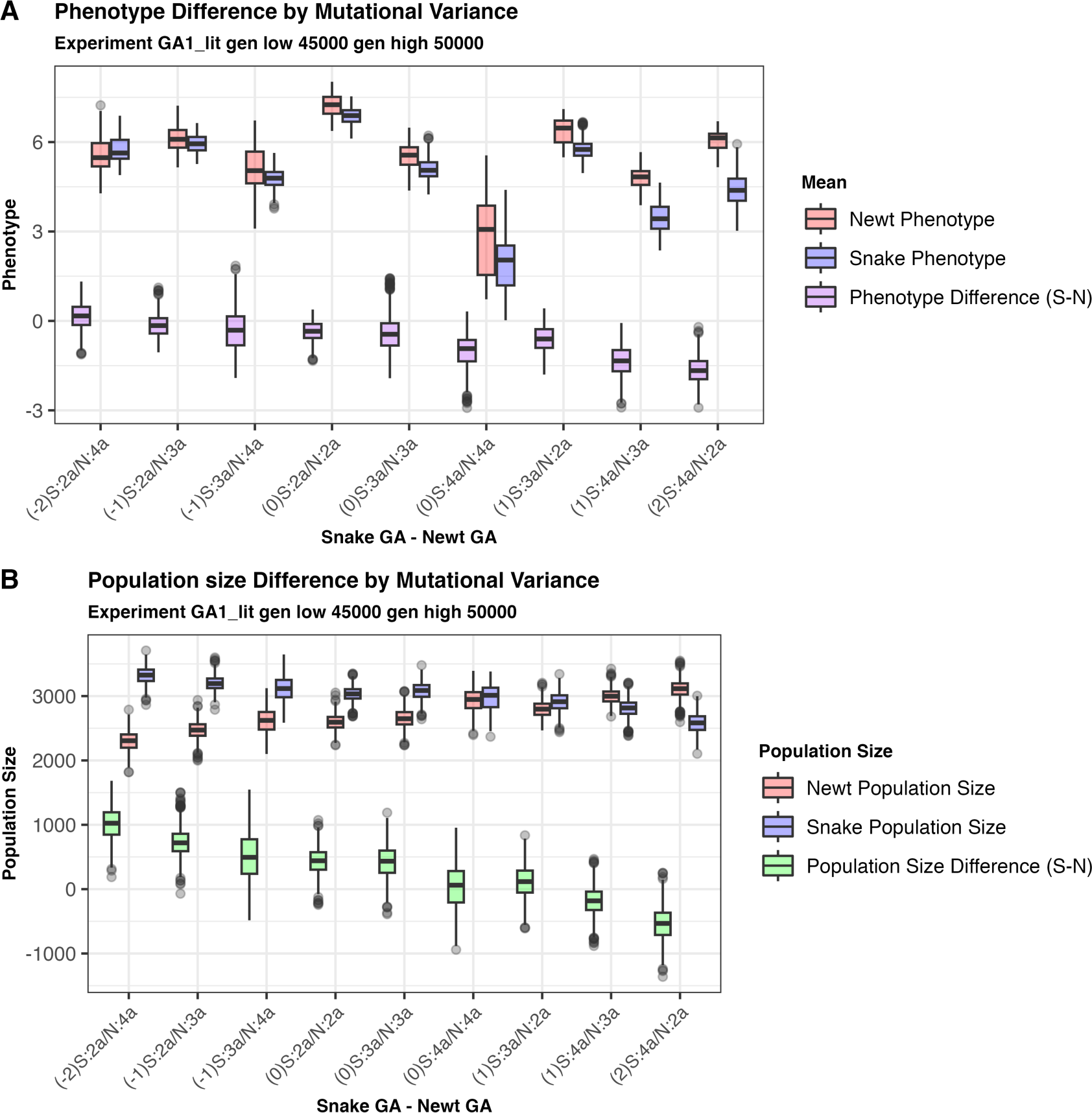
Distribution of (A) mean newt and snake phenotypes, and (B) newt and snake population sizes, and their differences (snake minus newt), from a regularly spaced set of generations between 45,000and 50,000. All combinations of genetic architecture from Experiment 1 are shown, except those containing the lowest mutational variance (1a). The combination of genetic architectures is shown on the *x*-axis labels, prepended with the difference of snake and newt log_10_(*V_M_*) values: for instance, the leftmost set of boxplots, labeled “(−2)S:2a/N:4a”, refers to simulations in which snakes have genetic architecture 2a, newts have genetic architecture 4a; and the snake’s genetic architecture has 100 times less mutational variance than does the newt’s.

Supplementary Figure S4 shows similar plots for Experiment 2, where the two species have different mutational variances but the same mutation rate, across four distinct mutation rates. In these simulations, although equilibrium phenotypes depended on the genetic architectures, in general the species with the larger mutational variance usually did *worse*, i.e., had a lower phenotype than the other species. (This is seen in Supplementary Figure S4 by the purple box plots moving down to the right.) However, when the two species had the same mutational variance, newts usually had higher phenotype than snakes.

On the other hand, Figure S5 shows that if both species have the same mutational variance, then differences in polygenicity do not have a strong effect on the outcome. (There are perhaps relatively minor differences in the equilibrium, but patterns are unclear.) In summary, we have good evidence for a strong effect of mutational variance, but not polygenicity, on which species is winning the coevolutionary arms race.

## 4 Discussion

In summary, we found that spatial heterogeneity in ecological factors was important for creating a spatial mosaic of correlated phenotypes. On the other hand, our simulations could not produce such coevolutionary mosaics in the absence of spatial heterogeneity, regardless of the genetic architecture. The genetic architecture did affect the dynamics of coevolution: the speed of coevolution depended primarily on mutational variance (in particular, if mutational variance was too low, the species would not coevolve). However, genetic architectures with the same mutational variance but different mutation rates and polygenicities had nearly identical coevolutionary dynamics.

One of our main goals was to identify conditions that would create the striking spatial correlations seen between the two species’ phenotypes in the wild. Even though our simulations had enough space for local adaptation to occur, we saw no such geographical correlations in simulations on uniform landscapes, regardless of genetic architecture. However, environmental gradients in various aspects of the underlying biology were sufficient to create geographical correlations of similar magnitude to that seen in the wild. This result, again, did not depend strongly on genetic architecture.

Genetic architecture affected the dynamics of coevolution: species with larger mutational variance could increase their phenotype faster (and so “win”, at least transiently). Furthermore, effects of genetic architecture on the final steady state seemed mostly attributable to mutational variance – genetic architectures with the same mutational variance but different polygenicities did not show large differences, at least over the fairly coarse scale we examined. Our results (e.g., Figure 7) suggest that there are indeed differences in the steady state achieved by different combinations of genetic architectures, but these differences are usually smaller than the generation-to-generation noise observed over the course of our simulations.

### Relationship to Other Models of Coevolution

Many previous theoretical papers also study coevolutionary dynamics. For instance, Nuismer, Thompson, and Gomulkiewicz (2000) study how coevolution might lead to clines in allele frequency using deterministic models in either discrete or continuous space in which each species carries a single biallelic locus that mediates the coevolutionary dynamics. Nuismer, Thompson, and Gomulkiewicz (2000) found many situations in which coevolution maintained spatial variation in allele frequencies, although their model did not include escalatory (i.e., arms race) antagonism. Nuismer, Ridenhour, and Oswald (2007) develop a quantiative model of trait coevolution in which – much like in our own – quatitative traits are additive subject to stabilizing selection, and the result of antagonistic inter-species interactions is determined by a logistic function of the difference in trait values (so that each species “tries to exceed” the other). Nuismer, Ridenhour, and Oswald (2007) deal with genetic architecture in three ways: by assuming fixed genetic variance and using a quantitative genetics model; by assuming weak selection and quasi-linkage equilibrium; and by doing deterministic numerical simulations in which genetic variance is due to a fixed number of explicitly represented loci evolving under strong selection. Nuismer, Gomulkiewicz, and Ridenhour (2010) also used a quantitative genetics model (i.e., of many small-effect loci but without explicitly representing the loci) in both analytical calculations and individual-based simulations of an island model. Nuismer, Gomulkiewicz, and Ridenhour (2010) found that conditions under which traits were correlated across space differed between across types of coevolutionary interactions. They also found differences between the analytical predictions and individual-based simulation results.

Our study looks at many of the same questions, using some of the same quantitative genetics tools. However, many aspects of models that are fixed in previous work (e.g., number of polymorphic loci or population size) are emergent properties of our simulations. We focus on using individual-based simulations to test how genetic architectures affect coevolutionary trajectories with explicit models of continuous space and genetic inheritance. It would not be possible to address some of our questions using previous models: for instance, we study a broad range of genetic architecture polygenicities, which would not be possible under a purely quantitative genetics model. Furthermore, our model explicitly represents ecological dynamics – so, for instance, local extinction/recolonization is possible, unlike under the fixed-population-density models of many previous papers. Additional points of realism of our simulations that make analysis of mathematical models difficult include recombination, fluctuating population sizes, and stochasticity. These are relatively straightforward to implement thanks to the simulation software, SLiM, especially in its’ newest release with improved support for interacting species (Haller and Messer 2023).

### What Does This Tell us About the Newts and Snakes?

Our results support the idea that the spatial patterns in toxicity and resistance observed in *Taricha* newts and *Thamnophis* garter snakes is a result of spatial heterogeneity in some ecologically important parameters, rather than simply spatial structure and resulting decoupling of local dynamics. It has been observed, for instance, that toxicity of one *Taricha* species is correlated with elevation in the Sierra Nevada (Reimche et al. 2020). There are many possible aspects that might vary across space; variation in both trait costliness and interaction rate had similar effects. Hanifin, Brodie, and Brodie (2008) presented data across a large area of the Pacific Northwest coast of North America, from Canada to Southern California, which spans a wide range of temperature, rainfall, and biodiversity. It is easy to imagine that, for instance, the costliness to toxin-resistant snakes of being less able to crawl quickly might vary with temperature, or that varying density of other species (e.g., toads or owls that prey on newts) might lead to spatial patterns in interaction rates (Toju and Sota 2006; Craig, Itami, and Horner 2007).

Feldman et al. (2010) hypothesized that snakes were “winning” the coevolutionary arms-race against newts due to the snakes’ genes of large effect. We found that mutational variance, rather than polygenicity, was the important determinant of which species was ahead (Figure 7). In fact, we found that in our simulations, newts – not snakes – had higher average phenotypes across most combinations of genetic architectures, although the effect was relatively small and somewhat hard to predict. Other scenarios are possible that would explain this situation: for instance, perhaps high phenotype values are more costly for newts than for snakes (in our simulations, we might set *c* smaller for newts than for snakes). Or, perhaps there are relatively hard constraints on the upper limit of tetrodotoxin production by newts, thus limiting the evolvability of the newt’s phenotype (which is particularly plausible if the toxin is produced by an environmentally-acquired bacteria (Vaelli et al. 2020)).

In summary, what have we learned about the newts and the snakes? Two major takeaways are that (a) the observed spatial correlations in phenotypes is entirely consistent with the proposed evolutionary story developed in the empirical literature, but that (b) existence of these spatial patterns does not strongly constrain the set of possible underlying genetic architectures.

### Spatial Patterns and Interactions

When evolution occurs across continuous space, evolutionary change and differentiation might lead to differences in different locations. However, we saw no such spatial patterns in simulations with a uniform environment. This might be due to the mixing action of migration, but even across a very large range, we still might not see substantial spatial patterns if evolution towards the same equilibrium was occurring at the same rate in all locations. (If ecological factors are heterogeneous, as in Figure 4, the equilibrium may differ across space.) In the real world, spatial patterns are not ubiquitous: for instance, Hoeksema and Forde (2008) did not find a relationship between spatial scale and degree of local adaptation.

Spatial patterns in biological systems are possible even without variation in the underlying environment. Even on a scale of meters, spatial patterns can spontaneously emerge from ecological dynamics, e.g., of vegetation in grasslands (Thompson and Daniels 2010; Sasaki 1997). Spatial cycling in reaction-diffusion equations is known to produce striking spatial patterns in certain circumstances (e.g., Murray 1982; Britton 1990; Schreiber and Killingback 2013). It could be, for instance, that local extinction-recolonization dynamics might play a role in the observed spatial patterns. This might work, for instance, by cycles of (a) newts evolve toxicity; (b) snakes evolve resistance; (c) snakes eat most of the newts; (d) snakes lose resistance; (e) newts recolonize from a nearby relatively snake-free environment. Alternatively, if snakes have an “all-or-nothing” genetic architecture of resistance and newts have relatively little genetic variance for toxicity at any time, then one might imagine a cycle of: (a) newts evolve toxicity; (b) snakes evolve resistance; (c) snakes are sufficiently resistant that variation in newt toxicity levels provides no advantage; (d) newts lose costly toxicity; (e) snakes lose resistance. Across space, these cycles could be decoupled between regions, and barriers to dispersal would make it more likely they are not synchronized, leading to spatial patterns. However, there is no evidence that such cycles occur in the real world. In this paper, we have not attempted to search for conditions that would produce such cycling; instead, our goal was to see whether, under reasonably plausible conditions, such dynamics might occur. It would, however, be interesting to investigate.

Our simulations highlighted the importance of understanding coevolutionary systems in a broader context: in our simulations, newts and snakes evolved in response not only to the selective pressure of coevolution, but also the heterogeneous environment. This point was also made by Yoder and Nuismer (2010), who suggested that heterogeneity in the environment was also necessary for coevolutionary dynamics to increase phenotypic diversity. Furthermore, similar coevolutionary mosaics could be produced by distinct genetic and environmental conditions, and so care must be taken in ascribing specific underlying mechanisms to patterns observed in the world (Craig and Itami 2021). This work highlights the importance of considering things outside of the focal species interaction (e.g., indirect interactions).

### Genetic Architecture

The underlying genetic mechanism of a trait determines how a trait can evolve and has implications on diversification and genetic variation (Hague, Feldman, et al. 2017). In our simulations, a species with sufficiently low mutational variance would never adapt, due to a lack of genetic variation. (For instance, if mutational variance for snakes was sufficiently low and newts were somewhat toxic, then rare resistance mutations in snakes might not increase snake phenotypes sufficiently on their own to meaningfully increase snake fitness.) Lack of genetic variation is thought to be a limiting factor in some potentially coevolutionary interactions (Hoeksema and Forde 2008).

Quantitative genetics models can predict how traits might change due to selective pressures (Lynch and Walsh 1998). These models typically depend only on genetic variance, without specifying the polygenicity of the underlying genetic architecture. We ran simulations under various architectures that ranged from simple (a few mutation of large effect) to complex (many mutations of minor effect), and although mutational variance indeed was the primary determining factor, polygenicity did affect steady state of the phenotype, although in complex ways. It would be interesting to model how deviations from the highly polygenic limit described by quantitative genetics affects results in practice.

Consistent with predictions of quantitative genetics, the speed of coevolution was mediated by the level of mutational variance created by the genetic architecture, rather than mutation rate or mutation effect size separately. The species that had a higher mutational variance often evolved and reached a steady state quicker. However, there were instances were the highest mutational variance limited a species’ ability to coevolve. This was potentially due to large effect mutations interacting with the costliness of the phenotype: a particularly large effect mutation might cause an individual to have a phenotype so high that it perished, thus slowing the speed of coevolution.

Perhaps counterintuitively, we found that higher mutational variance seemed to put a species at a coevolutionary *disadvantage* – see e.g., Figure 7. To understand this, consider what is happening in the simulations represented by the furthest-left boxplots of that figure. Here, newts have much higher mutational variance than snakes, and hence a much wider distribution of phenotypes: if this is sufficiently wide, then even if the mean newt phenotype is higher than the mean snake phenotype, there will still be many newts that can be eaten by the typical snake. Furthermore, snakes have a much higher population size than newts, so a given newt is more likely to encounter a snake than vice-versa, and so the selection pressure on newts to have a high phenotype is stronger than on snakes. A similar argument applies to the other end of the parameter range.

### Limitations and Continuing Questions

Our simulation was inspired by ecological interactions of real newts and snakes, but has a great many simplifications. For instance, newt and snake demographies (e.g., dispersal mechanism, birth rates, death rate, and age structure) were identical, despite *Taricha* newts living substantially longer (on average) than *Thamnophis* garter snakes in the wild. These aspects of life history impact the evolutionary trajectories of traits. Furthermore, many aspects of newt toxicity are unknown: it is thought that newt’s tetrodotoxin may be produced by a bacterium (as do other tetrodotoxin-producing species, Vaelli et al. 2020; Bucciarelli, Alsalek, et al. 2022b), but whether any such bacterium would be acquired from the environment or vertically transmitted is unknown. However, it seems likely that there would still be genetic variation related to the production and/or tolerance of the toxin.

In our simulation, if an interaction occurs between a newt and a snake, one perishes while the other survives. In the wild, a snake may attempt to eat a newt, fail, but both species survive (Reimche et al. 2020). Such interactions where both survive may lead to different results if trait plasticity is considered. It has been shown that a newt’s toxicity can increase and remain high for some time after the newt is be disturbed (Bucciarelli, Shaffer, et al. 2017). Therefore, a failed attack could leave a newt in a better position if more snakes were nearby. Relatively little is known about how often snakes and newts interact in the wild, and how much that varies across space or time.

Finally, it would be interesting to incorporate more specifics about the known genetic architecture of tetrodotoxin resistance. We did not pursue this direction in this paper because we were interested not only in conclusions about the real system, but also broader evolutionary questions about the role of genetic architecture in these sorts of coevolutionary dynamics. We have aimed more generally to capture important aspects of the dynamics without being distracted by unnecessary details. Modeling, particularly simulation-based exploration, walks a fine line between specificity (to faithfully represent an aspect of reality) and generality (so that results are generalizable). However, future work in this area could reflect new understandings of the system, as well as more realistically model differences between the species such as genome sizes and life cycles.

## Acknowledgments

We thank Heather Eisthen and Gary Bucciarelli for their fun conversations and helpful input. We also thank Gideon Bradburd and Bob Week for stimulating discussions about coevolution and early input on this project. This work was supported in part by NIH grants R56HG011395 and R01HG010774 from NIGMS to PR.

## Notes

https://github.com/Vcaudill/Coevolution

### Competing Interest Statement

The authors have declared no competing interest.

### Summary of Updates

Spelling and a correction to acknowledgments

