## Supplemental Figures for "Genetic architecture, spatial heterogeneity, and the coevolutionary arms race between newts and snakes"

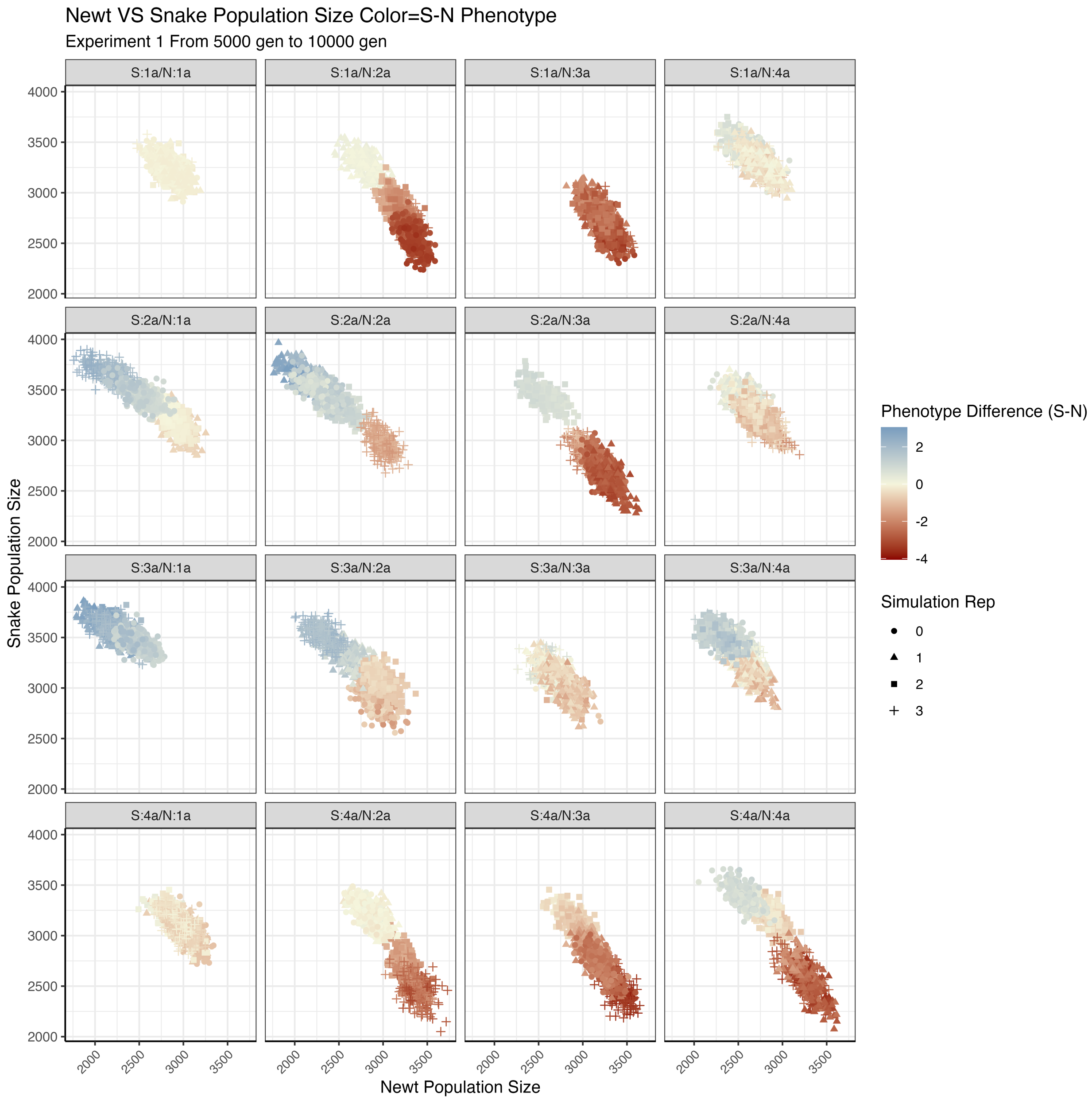


Figure S1: Newt and snake population sizes for the sixteen genetic architecture combinations of Experiment 1 over generations 5000 to 10000. Points are colored by the difference between average snake and average newt phenotypes. In each plot the population sizes at a evenly spaced set of time points are shown for each of the four replicates (replicate ID shown by point type). Newt and snake population sizes are generally negatively correlated: when newts have higher phenotypes (red points), newt populations tend to be larger and snake populations smaller. Note there is substantial variation between replicates, probably due to differing initial conditions.

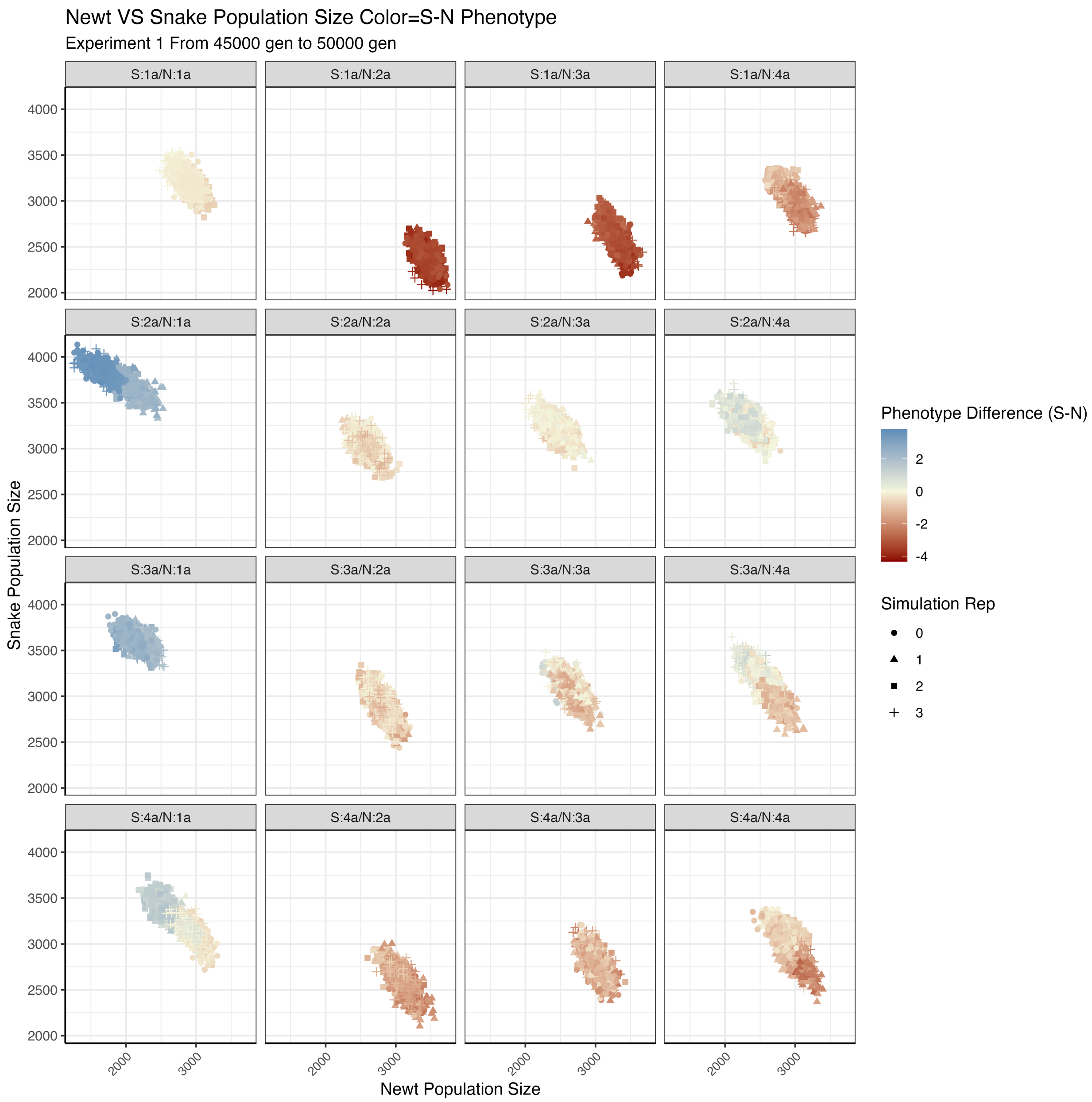


Figure S2: As in Figure S1, except that time points are between 45,000 and 50,000 generations (the end of the simulations). Note that there is less variation between replicates than in the earlier time points.

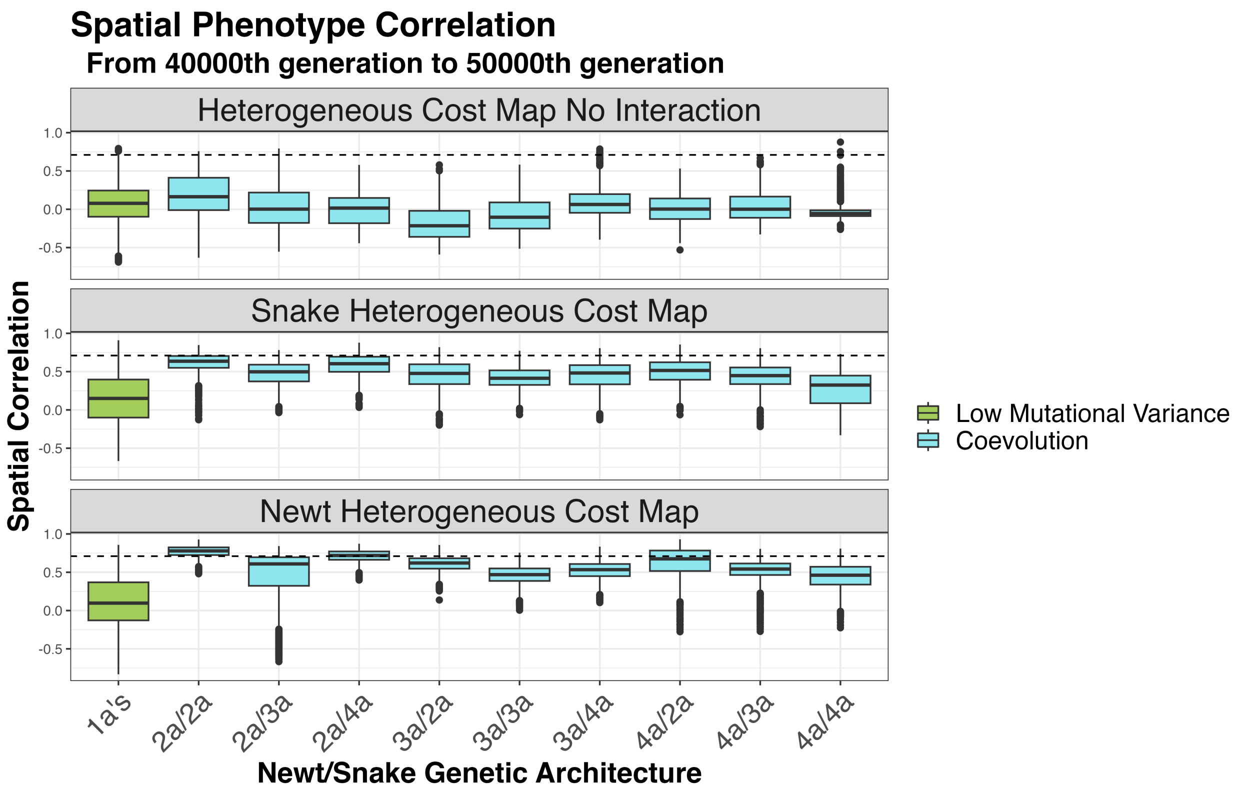


Figure S3: Distributions of spatial correlations between newt and snake phenotypes, across the combinations of genetic architectures of Experiment 1. Each boxplot shows the range of spatial correlations computed across an evenly spaced set of time points from 40,000 to 50,000 generations. **(top)** "Heterogeneous cost - no interaction" refers to simulations where costliness for both species varies across the landscape, but interaction outcome does not depend on phenotype (i.e., is just the result of a coin toss); **(middle)** "Heterogeneous cost - snake" refers to simulations that are as usual except that the costliness of the snake phenotype varies across the landscape (but not newt phenotypes); **(bottom)** "Heterogeneous cost - newt" refers to the converse situation, where newt phenotype costliness varies, but not snake.

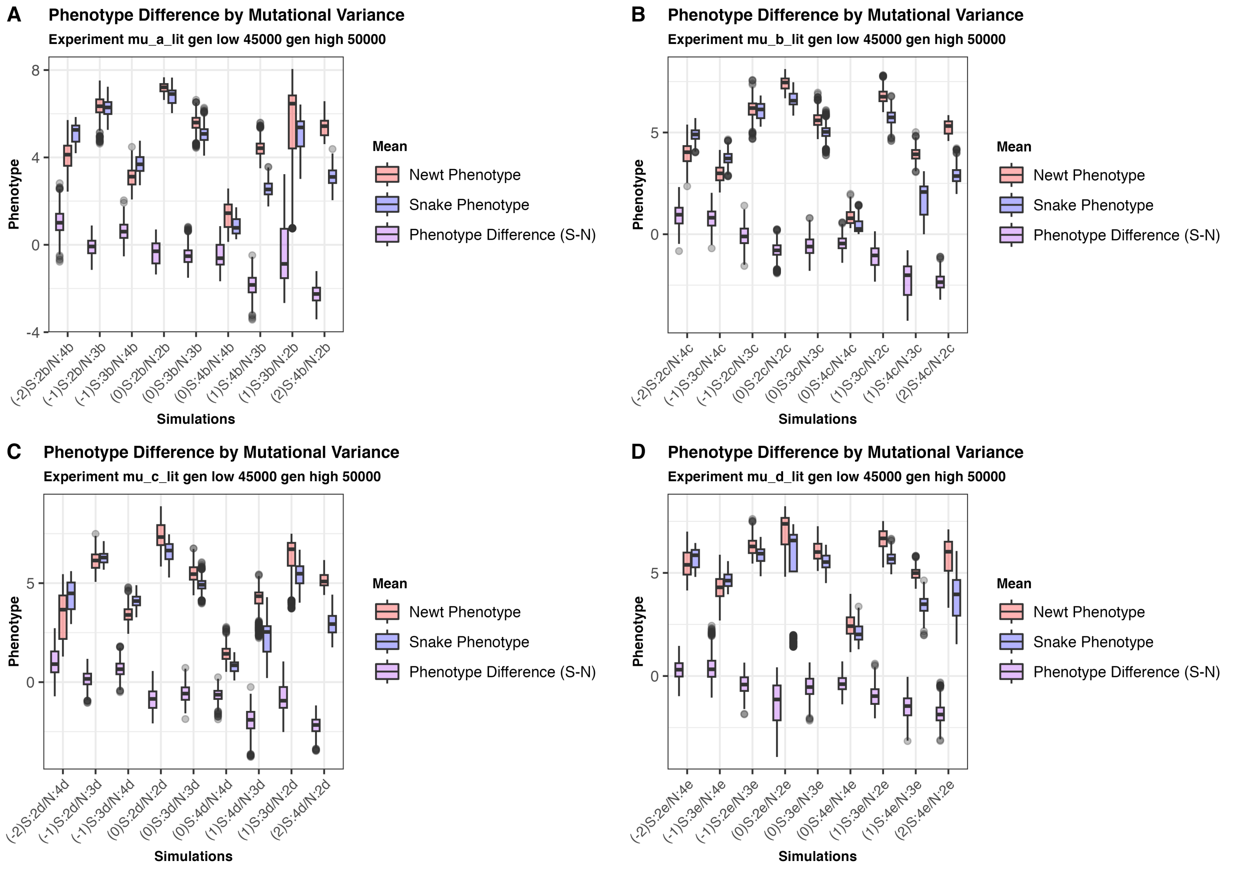


Figure S4: Distribution of mean newt and snake phenotypes and mean (snake minus newt) phenotype differences from a regularly spaced set of generations between 45,000 and 50,000. in **Experiment 2**, except for those simulations including a genetic architecture with the lowest mutational variance (1b, 1c, 1d, or 1e). (A) shows all nine combinations of mutational variances in the first shaded section of Experiment 2 of Table 1 (i.e., genetic architectures 2b, 3b, and 4b). The combination is shown on the $x$-axis labels, prepended with the difference of snake and newt $\log_{10}(V_M)$ values: for instance, the leftmost set of boxplots, labeled "(-2)S:2b/N:4b", refers to simulations in which snakes have genetic architecture 2b, newts have genetic architecture 4b; and the snake's genetic architecture has 100 times less mutational variance than does the newt's. (B, C, and D) show the same, but for combinations in the remaining three sections of Experiment 2 in Table 1. The arrangement is so that for boxplots on the left, snakes have lower mutational variance than newts, and on the right, newts have lower mutational variance than snakes.

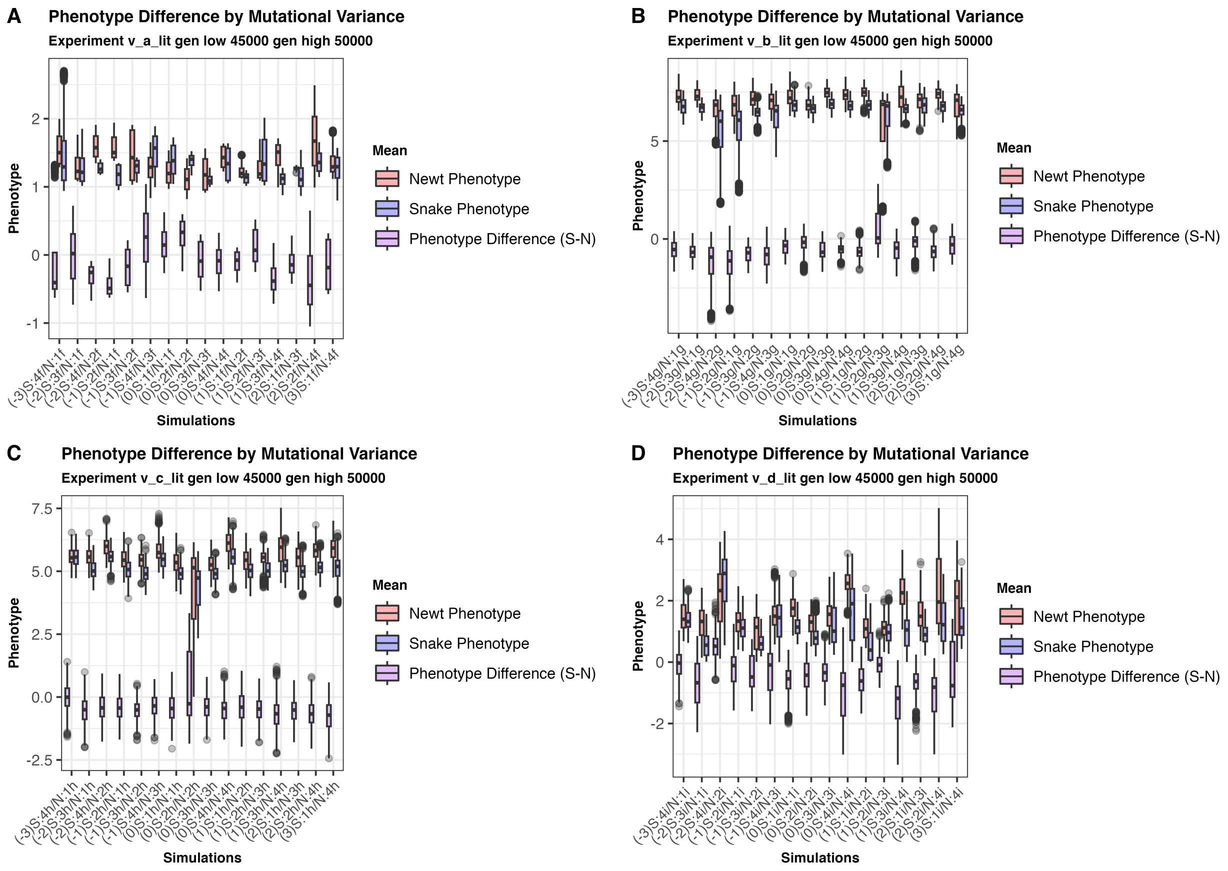


Figure S5: Mean newt and snake phenotypes (and mean differences), much as in Figure S4, but for **Experiment 3**. In this experiment, the different genetic architectures are grouped in Table 1 by mutation rate, so the boxplots are ordered by difference in $\log_{10}(\mu)$, so that on the left, snakes have lower mutation rate than newts, and on the right, newts have lower mutation rate than snakes.
